# Natural Variation of *OsLG3* Controls Drought Stress Tolerance in Rice by Inducing ROS Scavenging

**DOI:** 10.1101/228403

**Authors:** Haiyan Xiong, Jianping Yu, Jinjie Li, Xin Wang, Pengli Liu, Hongliang Zhang, Yan Zhao, Zhigang Yin, Yang Li, Yan Guo, Binying Fu, Wensheng Wang, Zhikang Li, Jauhar Ali, Zichao Li

## Abstract

**Background:** Improving performance of rice under drought stress has potential to significant impact on rice productivity. Previously we reported that *OsLG3* positively control rice grain length and yield.

**Results:** In this study, we found that *OsLG3* was more strongly expressed in upland rice compared to lowland rice under drought stress condition. Candidate gene association analysis showed that the natural variation in *OsLG3* was associated with tolerance to water deficit stress in germinating rice seeds. Transgenic rice with enhanced *OsLG3* expression exhibited improved tolerance to drought and that is most likely due to enhanced ROS scavenging efficiency. Phylogenetic analysis and pedigree records indicated that the tolerant allele of *OsLG3* has potential to improve drought tolerance of *japonica* rice.

**Conclusions:** Collectively, our work revealed that the natural variation of *OsLG3* contributes to rice drought tolerance and the elite allele of *OsLG3* is a promising genetic resource for the development of drought-tolerant and high-yield rice varieties.

## Background

Agriculture experience more than 50% of average yield losses worldwide due to abiotic stress, especially drought [1, 2]. Rice (*Oryza sativa* L.), one of the most important staple food for over half the world’s population, requires high inputs of water during growth, which results in a number of production challenges due to water shortages and inadequate rainfall during the rice growing season [3]. In the face of these challenges, enhanced performance of rice under drought stress has the potential to significantly improve rice productivity.

Plants have evolved complex signaling pathways that enable them to respond and adapt to unfavorable environmental conditions through morphological, physiological, and biochemical changes [4-6]. The generation of reactive oxygen species (ROS) is a key process in plant responsiveness to various biotic and abiotic stresses. ROS are important signal transduction molecules, but also toxic by-products of stress metabolism, depending on their overall cellular amount [7]. At low to moderate concentrations, ROS are likely to function as secondary messengers in stress-signaling pathways, triggering stress defense/adaptation reactions. However, when ROS levels reach above a certain threshold, they can trigger progressive oxidative damage resulting in retarded growth and eventually cell death [8]. Plants have developed flexible ROS scavenging and regulation pathways to maintain homeostasis and avoid over-accumulation of ROS in cells. There are some evidences supporting that overproduction of ROS-scavenging related genes and synthesis of various functional proteins such as superoxide dismutase (SOD), peroxidase (POD) and catalase (CAT) could increase tolerance to oxidative stress. For example, overexpression of *SNAC3* (Stress-responsive NAM, ATAF1/2, and CUC2 gene 3), increases rice drought and heat tolerance by modulating ROS homeostasis through regulating the expression of genes participated in ROS-scavenging [9]. Overexpression of an Ethylene-responsive factor (ERF) family gene, *SERF1* (Salt-Responsive ERF1), improved salinity tolerance in rice, mainly due to the regulation of ROS-dependent signaling during the initial phase of salt stress [10]. Overexpressing of *JERF3* (Jasmonate and Ethylene-Responsive Factor 3) improved the expression of genes involved in ROS-scavenging and enhanced tolerance to drought, freezing, and salt in tobacco [11]. Moreover, overexpression of the mitogen-activated protein kinase kinase kinase (MAPKKK) gene *DSM1* (Drought-hypersensitive Mutant 1) in rice, increased drought stress tolerance by regulating ROS scavenging. Conversely, deficiency in *DSM1* resulted in a decrease in ROS scavenging and increased drought hypersensitivity [12].

Transcription factors (TFs) play central roles in the regulation of gene expression in the stress signaling and adaptation networks [13, 14]. One set of these is the APETALA2/Ethylene Responsive Factor (AP2/ERF) superfamily. It has been implicated that the ERF proteins play diverse roles in cellular processes involving flower development, spikelet meristem determinacy, floral meristem, plant growth, pathogens and abiotic stress tolerance [15-23]. There are emerging evidences to support that ERF proteins are involved in response and adaptation to drought stress. For instance, transgenic rice lines overexpression ERF TFs including SUB1A, *OsEREBP1*, *AP37*, *AP59*, *HYR* and *OsERF71*, all showed strong resistance to drought stress [24-28]. Other ERF genes including *HARDY*, *TRANSLUCENT GREEN* (*TG*) [29] and DREB genes [30, 31] from *Arabidopsis*, *TSRF1* [32] from tomato, *TaERF3* [33] from wheat (*Triticum aestivum*), *GmERF3* [34] from soybean, *JERF3* [11] from tobacco and *SodERF3* [35] from sugarcane (*Saccharum officinarum*), have also been found to be involved in responses to water deficit stress condition. Overall, these findings suggested that ERF TFs offer the potential for engineering crops in a way that makes them more efficient under drought stress condition.

Linkage disequilibrium (LD)-based association mapping has been proven to be a powerful tool for dissecting complex agronomic traits and identifying alleles that can contribute to crop improvement [36-39]. Candidate gene association analysis, an effective method to validate targets, has become easier and cheaper with the advances in next-generation sequencing (NGS) technology facilitating the discovery and detection of single nucleotide polymorphisms (SNPs) [40]. This strategy has been used successfully in the genetic dissection of allelic diversity of genes controlling fatty acid content, kernel size, ABA content, α-tocopherol and β-carotene content and aluminum tolerance in maize and rice [41, 42]. There have also been reports of association studies on crop drought tolerance. For instance, Y Lu, S Zhang, T Shah, C Xie, Z Hao, X Li, M Farkhari, JM Ribaut, M Cao, T Rong, et al. [43] and Y Xue, ML Warburton, M Sawkins, X Zhang, T Setter, Y Xu, P Grudloyma, J Gethi, J-M Ribaut, W Li, et al. [44] identified some QTLs underlying drought tolerance in maize by genome-wide association analysis. S Liu, X Wang, H Wang, H Xin, X Yang, J Yan, J Li, LS Tran, K Shinozaki, K Yamaguchi-Shinozaki, et al. [45] found that DNA polymorphisms in the promoter region of *ZmDREB2.7* were associated with maize drought tolerance. Analysis of the association found that an 82 bp insertion in *ZmNAC111* and 366 bp insertion in *ZmVPP1* affected drought tolerance in Maize [46, 47]. Recently, an association study of 136 wild and four cultivated rice accessions identified three coding SNPs and one haplotype in a DREB (Dehydration Responsive Element Binding) transcription factor (TF), *OsDREB1F*, that are potentially associated with drought tolerance [48], and nine candidate SNPs were identified by association mapping of the ratio of deep rooting in rice [49]. However, a role for these candidate genes or their causative variations in enhanced drought tolerance remains to be verified experimentally.

Here, we characterize the role of an ERF family TF, *OsLG3* (LOC_Os03g08470), in rice drought tolerance. *OsLG3* is located at the same locus as *OsERF62* [23] and *OsRAF*(a Root Abundant Factor gene in *Oryza sativa*) [50]. In our previous work, we demonstrated that *OsLG3* plays a positive role in rice grain length without affecting the grain quality [51]. In this study, we identified the natural variation in the promoter region of *OsLG3* was associated with tolerance to drought stress among different rice accessions. Overexpression of *OsLG3* in transgenic lines resulted in increased drought tolerance via regulating ROS homeostasis. *OsLG3* function as a pleiotropic gene which can contribute to rice grain length and drought stress tolerance together. These data provide insights that the elite allele of *OsLG3* is a promising genetic resource for the genetic improvement of rice drought tolerance and yield.

## Results

### *OsLG3* is associated with drought stress tolerance in rice

In our previous cDNA microarray experiment, comparison between upland rice (UR) (IRAT109 and Haogelao, drought-resistant *japonica* rice) and lowland rice (LR) (Nipponbare and Yuefu, drought-sensitive *japonica* rice) varieties under well-watered and water deficit conditions showed that *OsLG3* expression was induced to a significantly greater extent by water deficit stress in UR than LR [52]. To confirm this we analyzed the expression level of *OsLG3* between IRAT109 and Nipponbare under increasingly severe water deficit conditions using Quantitative real-time PCR (qRT-PCR). The data confirmed that *OsLG3* is more highly expressed in IRAT109 than Nipponbare under well-watered conditions and that *OsLG3* expression is strongly induced by drought in IRAT109 but not Nipponbare (**Figure. 1a**). These results suggested that changes in *OsLG3* expression may be involved in response to drought stresses.

**Figure 1.**
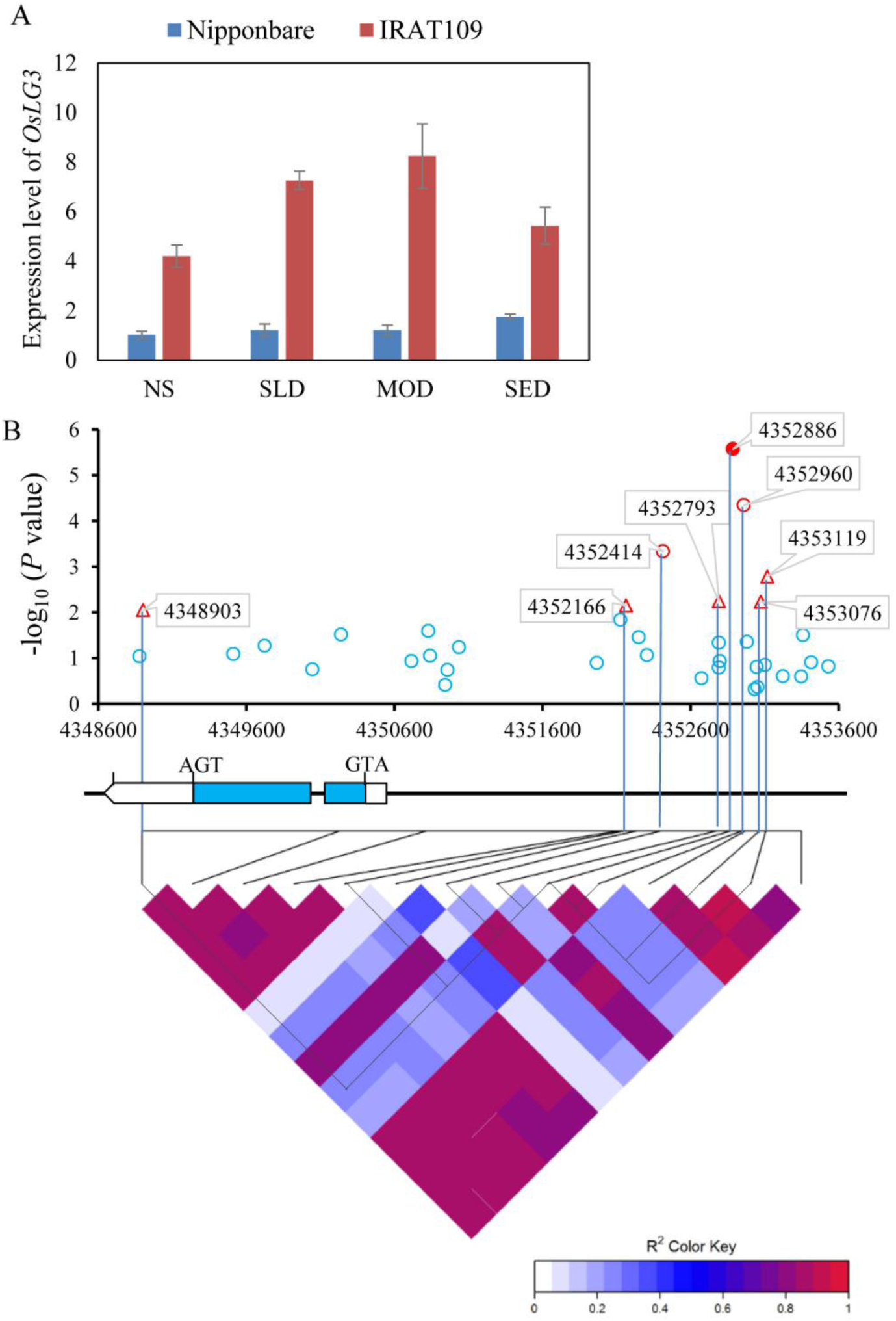
*OsLG3* is associated with drought tolerance. **(a)** qRT-PCR analysis of *OsLG3* in Nipponbare and IRAT109 under different soil drought stress levels. NS, No stress; SLD, Slight drought; MOD, Moderate drought; SED, Severe drought. **(b)** Analysis of the association between pairwise LD of DNA polymorphisms in the *OsLG3* gene and water deficit tolerance. Schematic of OsLG3 is shown on the X-axis and the significance of each variation associated with seedling RGR (ratio of germination rates under 15%PEG condition to germination rates under water conditions) is shown on the Y-axis. The SNPs with significant variation (*P* < 1.0 × 10^-2^) between genotypes are connected to the pairwise LD diagram with a solid line. Black dots in the pairwise LD diagram highlight the strong LD of SNP_4352886 (filled red circle, *P* = 2.66 × 10^-6^) and two significant variations, SNP_4352414 and SNP_4352960 (open red circles, *P* < 1.0 × 10^-3^). The SNP_4348903, SNP_4352166, SNP_4352793, SNP_4353076 and SNP_4353119, which are marginally significant (*P* < 1.0 × 10^-2^) are denoted by red triangles.

We conducted a candidate gene association analysis to investigate if natural variation in *OsLG3* is associated with rice drought tolerance. This approach utilized a mini core collection (MCC) panel [53] (**Additional file 7: Table S1**) of 173 varieties that have undergone deep sequencing to a 14.9× average depth (http://www.rmbreeding.cn/Index/). Seeds from MCC lines were germinated on water or in the presence of 15% polyethylene glycol (PEG) to simulate drought stress. The relative germination rate (RGR, ratio of germination rates under 15%PEG condition to germination rates under water condition) of each line was calculated after five days. We identified significant variation in water deficit stress tolerance between the different varieties (**Additional file 1: Figure S1** and **Additional file 7: Table S1**). A total of 97 SNPs within the *OsLG3* locus from these accessions were identified. To reduce the incidence of false positives, we performed general linear model (GLM) that controls population structure (Q matrix) to identify significant genotypic and phenotypic associations. The association analysis detected three significant SNPs (*P* < 1.0 × 10^-3^) (SNP_4352414, SNP_4352886, SNP_4352960) located within the promoter region of *OsLG3* (**Figure 1b**). SNP_4352886, located 2449 bp upstream from the start codon of *OsLG3*, showed greatest significant association with RGR (*P* = 2.66 × 10^-6^, **Figure 1b**), contributed 13.9% of the phenotypic variation in the MCC population. SNP_4352886 was in strong LD with two other variations (SNP_4352414, SNP_4352960) in the promoter (r^2^ **≥** 0.8), but not with the SNP_4348903, SNP_4352166, SNP_4352793, SNP_4353076 and SNP_4353119, which were identified as marginally significant (*P* < 1.0 × 10^-2^) (**Figure 1b**). SNPs identified within the coding region of *OsLG3* were not significantly associated with the RGR trait. Based on results above, we conclude that the nucleotide polymorphisms in the promoter of *OsLG3* are associated with differential germination rates under water deficit.

### OsLG3 is an ERF family transcription activator that functions as a homodimer

*OsLG3* encodes a putative protein with 334 amino acids. Amino acids 110-159 contain a typical AP2 domain, including 11 putative DNA-binding sites, implying a strong DNA binding capacity, and one putative nuclear localization signal (NLS) from amino acids 95-121 (**Additional file 2: Figure S2a, b**). Phylogenetic analysis comparing *OsLG3* with known ERF TFs [23] indicated that OsLG3 belongs to group VII of the ERF subfamily and is closely related to OsERF71 [27, 54], OsEREBP1 [25] and OsBIERF1[55], which have been reported to be involved in stress response (**Figure 2**).

**Figure 2.**
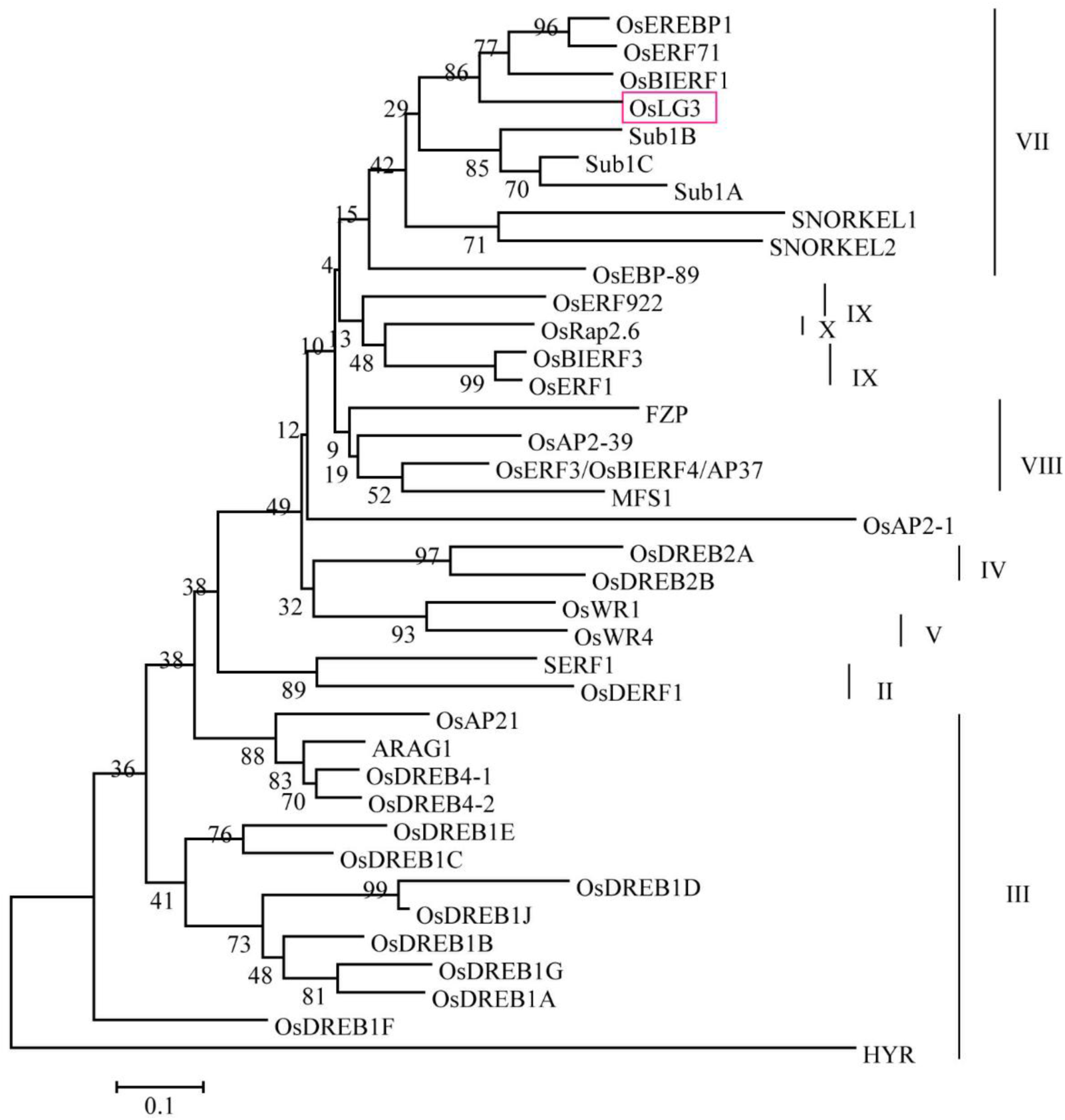
Phylogenetic tree of OsLG3 homologues. Neighbour-Joining phylogenetic analysis of OsLG3 protein sequence in the context of other characterized AP2/ERF proteins from rice. The phylogenetic tree was constructed using the ClustalW and MEGA programs. Tree topology with bootstrap support is based on a percentage of 1000 replicates. Accession numbers are as follows:ARAG1, LOC_Os02g43970; OsAP21, LOC_Os01g10370; OsDREB4-1, LOC_Os02g43940; OsDREB4-2, LOC_Os04g46400; OsDREB2A, LOC_Os01g07120; OsDREB2B, LOC_Os05g27930; OsBIERF3, LOC_Os02g43790; OsERF1, LOC_Os04g46220; OsERF922, LOC_Os01g54890; OsWR1, LOC_Os02g10760, OsWR4, LOC_Os06g08340; SNORKEL1, AB510478; SNORKEL2, AB510479; OsBIERF1, LOC_Os09g26420; OsEBP-89, LOC_Os03g08460; OsEREBP1, LOC_Os02g54160; OsLG3, LOC_Os03g08470; OsERF71, LOC_Os06g09390; Sub1A, DQ011598; Sub1C, LOC_Os09g11480; Sub1B, LOC_Os09g11460; MFS1, LOC_Os05g41760; OsAP2-39, LOC_Os04g52090; OsERF3/OsBIERF4/AP37, LOC_Os01g58420; FZP, LOC_Os07g47330; OsRap2.6, LOC_Os04g32620; OsAP2-1, LOC_Os11g03540; OsDREB1D, LOC_Os06g06970.

Transactivation activity assays, in which the DNA-binding domain (GAL4-BD) of GAL4 was fused to either the full length CDS of OsLG3 or a series of shortened fragments created by deletions from both the N- and C-termini, indicated that OsLG3 is capable of transcriptional activation, and that the C-terminal region (amino acids 213-334) is required for this (**Figure 3a**). We also tested dimerization of OsLG3 protein *in vivo* using a yeast two-hybrid system. To avoid the interference caused by self-transactivation activity, we used the protein fragment BD-dC2 (amino acids 1-218) as bait, which lacks the transactivation region. The constructs AD-OsLG3 and BD-dC2 were co-transformed into yeast strain AH109 to test for a homodimer interaction. AD and BD, AD-OsLG3 and BD were also co-transformed as negative controls for endogenous transactivation activity. Of these combinations, only AD-OsLG3 and BD-dC2 co-transformants could grow well in the SD/-Trp-Leu-Ade-His/X-α-gal medium (**Figure 3b**). These results indicate that OsLG3 potentially functions as a homodimer in rice. To determine the subcellular localization of OsLG3, *Nicotiana benthamiana* leaves were infiltrated with *Agrobacterium tumefaciens* (strain EH105) containing 35S:OsLG3-GFP and 35S::GFP, respectively. Confocal imaging analyses showed nuclear-localized fluorescence in 35S:OsLG3-GFP transformed cells, whilst fluorescence from free GFP was distributed throughout the whole cell (**Figure 3c**). This indicates that OsLG3 is a nuclear-localized protein.

**Figure 3.**
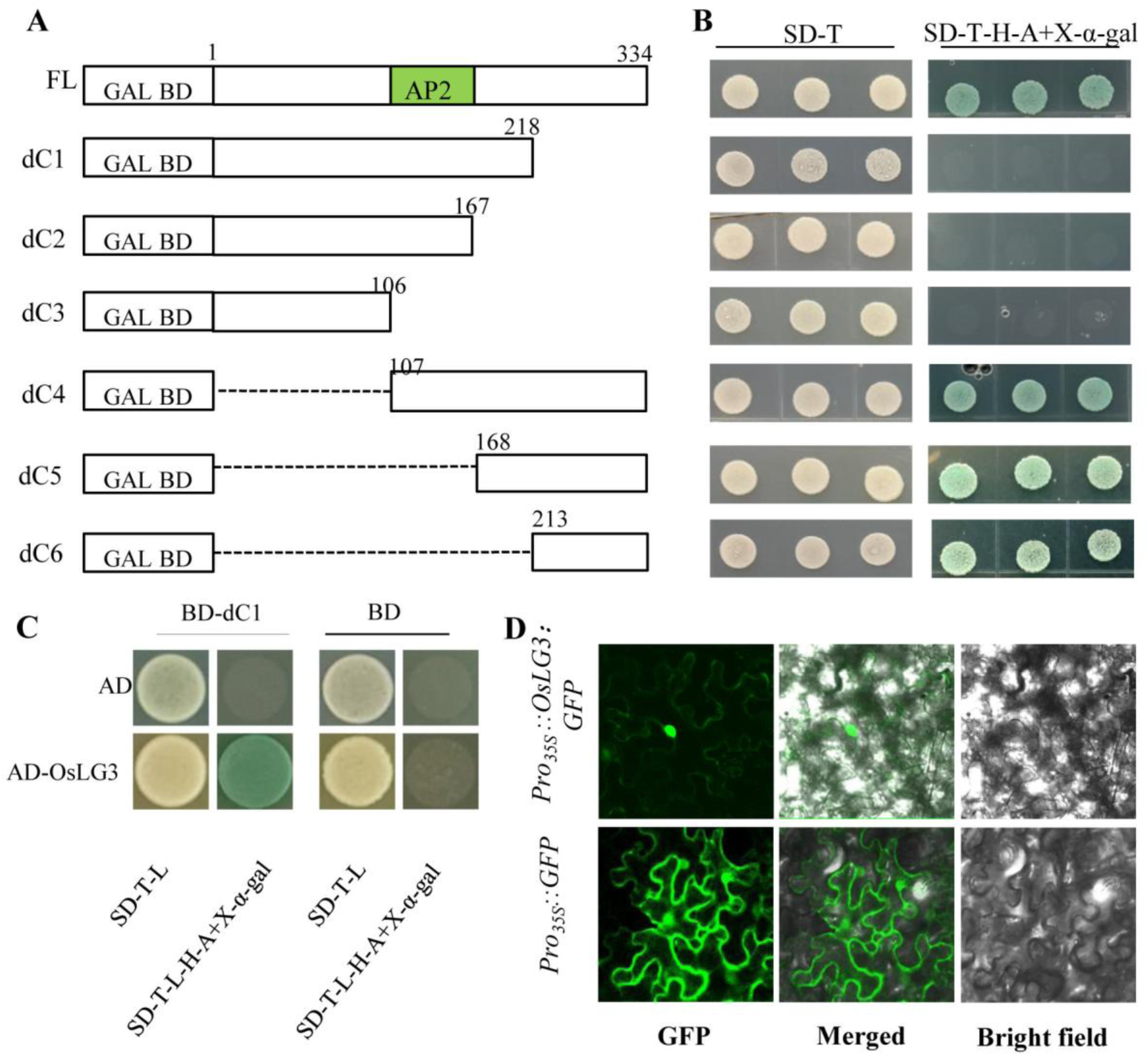
Transactivation assay and Subcellular localization of OsLG3. **(a)**Transactivation assay of OsLG3 fragments. Fusion proteins of the GAL4 DNA-binding domain and different portions of OsLG3 were expressed in yeast strain AH109 and streaked on control plate (SD/-Trp) or selective plate (SD/-Trp-Ade-His/X-α-gal), respectively. FL indicates the full-length CDS of OsLG3; dC1 to dC6 indicate the mutated forms of OsLG3 (nucleotide positions are labeled in the diagrams), respectively. **(b)** Dimerization analysis. BD with AD empty vector, AD empty with BD-dC2, AD-OsLG3 with BD empty and AD-OsLG3 with BD-dC2, were co-transformed in to yeast strain AH109 yeast cells and streaked on control plate (SD/-Trp-Leu) or selective plate (SD/-Trp-Leu-Ade-His/X-α-gal), respectively. The plates were incubated for 3 days. **(c)** Subcellular localization of OsLG3 in tobacco epidermal cells. The upper panels show the localization of OsLG3-GFP in onion cells in a transient assay, while bottom panels show the localization of GFP as a control.

### Expression profile of *OsLG3* under different stress treatments and in different plant tissues

The expression profile of *OsLG3* in response to abiotic stresses and hormones in IRAT109 was investigated using qRT-PCR. The transcript level of *OsLG3* was significantly induced under dehydration, polyethylene glycol (PEG), hydrogen peroxide (H_2_O_2_), NaCl, abscisic acid (ABA), ethylene (ETH) and gibberellin (GA) treatments but remained unchanged under cold treatment (**Figure 4a**). To determine the spatial-temporal expression of *OsLG3* under normal growth conditions, we isolated total RNA in eight representative tissues (root, stem, sheath at seedling stage and root, stem, sheath, leaf, panicle at reproductive stage) from IRAT109, and performed qRT-PCR analysis. *OsLG3* was expressed in all of the tissues tested, and showed higher level in roots compared to other tissues (**Figure 4b**).

**Figure 4.**
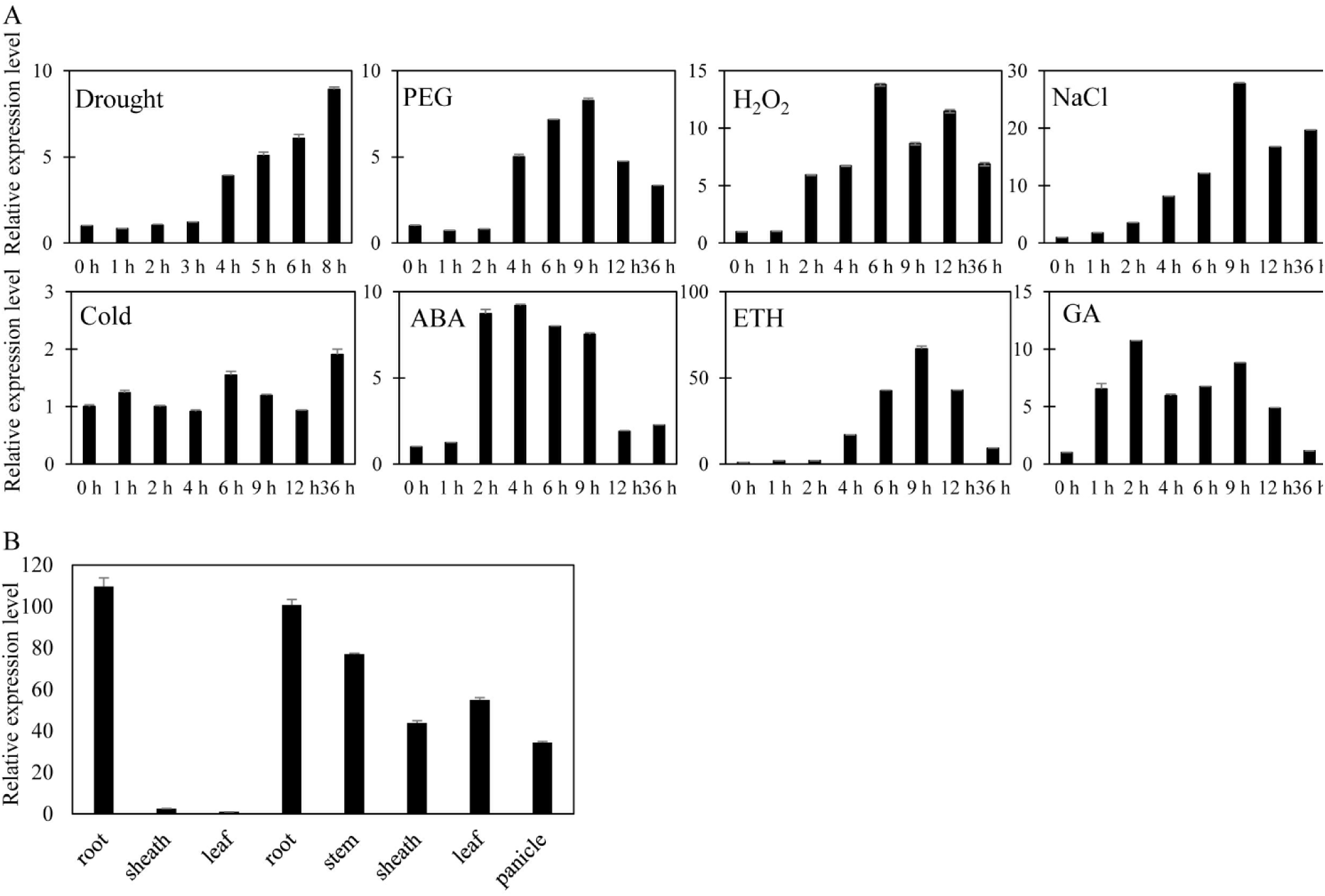
Expression analysis of *OsLG3.* **(a)** Expression level of *OsLG3* under various abiotic stresses and hormone treatments in IRAT109. Three-week-old seedlings were subjected to dehydration, NaCl (200 mmol), PEG 6000 (20%, w/v), cold (4°C), H_2_O_2_ (1 mM), ABA (100 μM), ETH (100 μM) and GA (100 μM) treatment. The relative expression level of *OsLG3* was detected by qRT-PCR at the indicated times. Error bars indicated the SE based on three replicates. **(b)** Detection of *OsLG3* expression in various tissues and organs of IRAT109 using qRT-PCR. Three-week-old seedlings were used to harvest the samples of the root, sheath and leaf at seedling stage. Plants before heading stage were used to harvest the samples of root, stem, sheath, leaf and panicle at the reproductive growth stage. Error bars indicate the SE based on three technical replicates.

### Overexpression of *OsLG3* enhances drought stress tolerance in rice

To elucidate the biological functions of *OsLG3* during the stress response, 10 independent transgenic lines in which *OsLG3* was overexpressed under the control of 35S promoter *(OsLG3*-OE) and 14 independent transgenic rice lines in which endogenous *OsLG3* expression level was suppressed by RNA interference (*OsLG3*-RNAi) were generated. qRT-PCR was carried out to quantify *OsLG3* expression level in each transgenic line. Two independent *OsLG3*-OE transgenic lines (OE4, OE7) with highest expression level (**Additional file 3: Figure S3**) and two *OsLG3*-RNAi lines (RI6, RI10) with lowest expression level (**Additional file 4: Figure S4**) of *OsLG3* were selected for further stress analysis.

Under dehydration treatment using 20%PEG for 3 days, *OsLG3*-OE lines showed greater resistance than WT plants (**Figure 5a**). Almost 44-81% of *OsLG3*-OE plants survived, while only 8-15% of the WT plants survived under this treatment (**Figure 5c**). In contrast, when four weeks old WT and *OsLG3*-RNAi plants were stressed by a slightly less severe dehydration stress (20% PEG for 2.5 days) the relative survival of *OsLG3*-RNAi lines (3-11%) was lower than that of WT (24-42 %) (**Figure 5b, d**).

**Figure 5.**
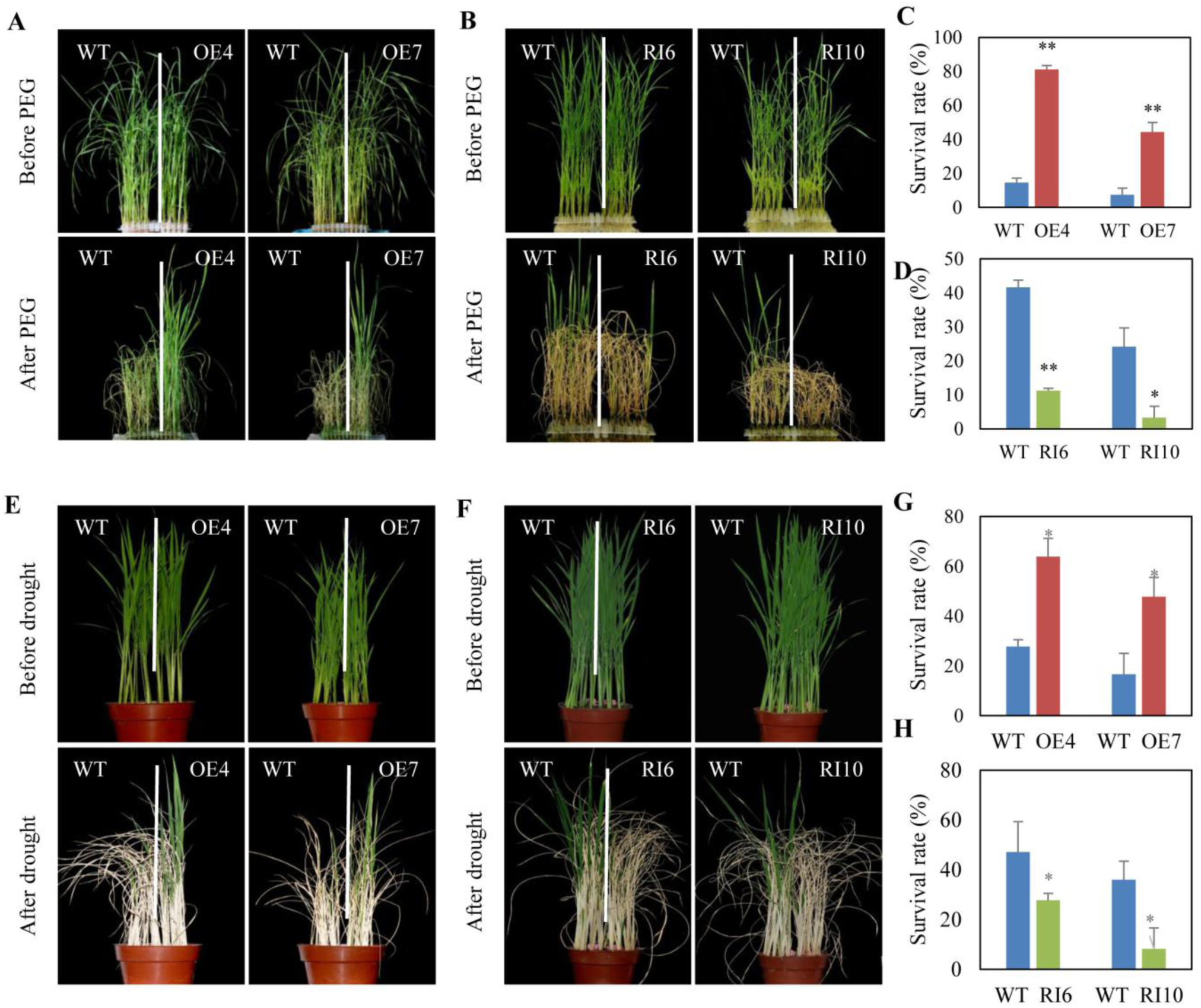
*OsLG3* increases rice survival under severe drought stress. **(a)** Physiological dehydration stress tolerance assay; *OsLG3-OE* plants subjected to 20% PEG for 3 days before being allowed to recover for 10 days. **(b)** Physiological dehydration stress tolerance assay of *OsLG3*-RNAi plants subjected to 20% PEG for 2.5 days before being allowed to recover for 10 days. **(c)** and **(d)**, Survival rates of transgenic and WT plants testing in **(a)** and **(b)**. Values are means ± SE (n = 3). **(e)** *OsLG3*-OE and WT plants were subjected to severe drought stress without water for 7 days and then recovered for 10 days. **(f)** *OsLG3*-RNAi and WT plants were subjected to moderate drought stress without water for 6 days and then recovered for 10 days. **(g)** and **(h)**, Survival rates of transgenic and WT plants testing in **(c)** and **(d)**. Values are means ± SE (n = 3). * and ** indicate significant difference at *P* < 0.05 and *P* < 0.01, respectively.

Under severe drought soil drought stress condition (stop watering for 7 days), the *OsLG3*-OE lines showed an improved survival rate compared to WT plants. Almost 48-64% of *OsLG3*-OE survived, whereas only 17-28% of WT plants survived this treatment (**Figure 5e, g**). In contrast, when WT and *OsLG3*-RNAi plants were treated moderate drought (stop watering for 6 days), 36-47% of the WT plants had recovered 10 days after watering was restored, but only 8-28% of the RNAi plants recovered (**Figure 5f, h**).

Interestingly, we found that the leaves of *OsLG3*-OE plants showed a slower rate of water loss compared to those of WT and *OsLG3*-RNAi lines under dehydration condition (**Additional file 5: Figure S5a**), indicating a role for *OsLG3* in reducing water loss, especially under water deficit conditions. Expression levels of some characterized stress response genes like *OsLEA3*, *OsAP37*, *SNAC1*, *RAB16C*, *RAB21* and *OsbZIP73* were monitored in the WT, *OsLG3*-OE and RNAi lines under well-watered and soil drought conditions. As shown in **Additional file 5: Figure S5b**, these genes showed significantly higher levels under soil drought stress in *OsLG3-OE* plants when compared to WT and RNAi lines.

We also analyzed the growth of *OsLG3*-OE and -RNAi plants under NaCl and mannitol treatment to induce high salinity and osmotic stress, respectively. Under NaCl treatment, OE lines showed significantly less suppression of relative shoot growth (shoot length under stress conditions to shoot length under normal conditions) than both WT and RNAi lines, and less suppression of relative fresh weight (fresh weight under stress conditions to fresh weight under normal conditions) than WT and RNAi lines (**Figure 6a, c**). Under mannitol treatment, the relative shoot growth of OE plants was significantly higher than that of both WT and RNAi lines, and the relative fresh weight in OE lines was higher than that of WT and RNAi lines (**Figure 6b, d**). These results show that changes in the expression level of *OsLG3* have a significant effect on the drought tolerance in rice.

**Figure 6.**
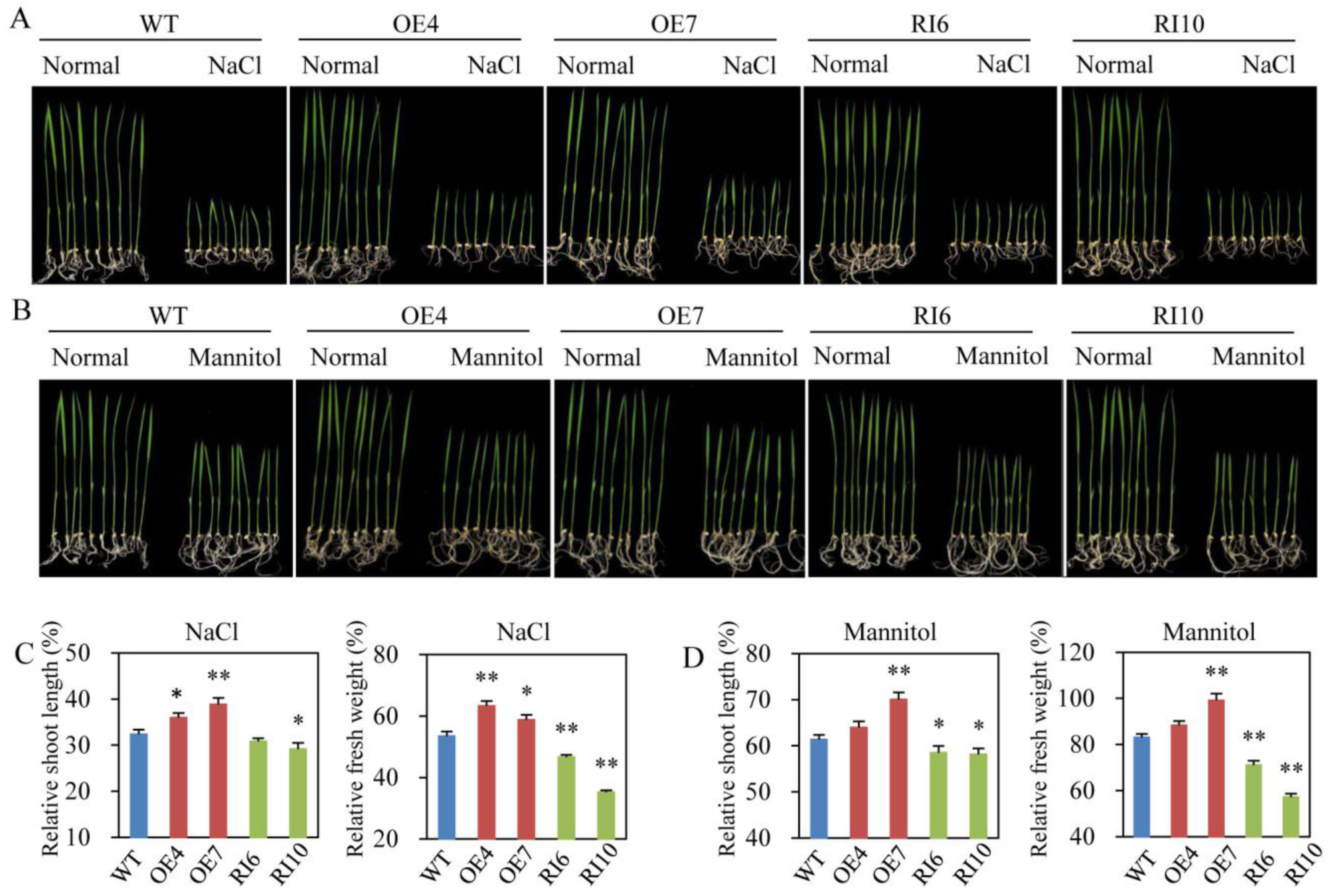
Growth of *OsLG3* transgenic plants under high salinity and osmotic stress conditions. Ten day old seedlings of WT, *OsLG3*-OE and RNAi plants grown in ½ MS medium containing **(a)** 150 mmol/L NaCl or **(b)** 200 mmol/L mannitol. The relative shoot length **(c)** and relative fresh weight **(d)** of the transgenic and WT plants from **(a)** and **(b)** were compared. Values are means ± SE (n = 10). * and ** indicate significant differences at *P* < 0.05 and *P* < 0.01, respectively.

### Global gene expression analysis revealed alteration in the expression of stress-related genes and ROS-scavenging related genes

Preliminary evidence for altered expression of stress-related genes in *OsLG3* -OE and RNAi lines led us to investigate the pathways regulated by *OsLG3* through transcriptome analysis. RNA samples were isolated from the leaves of ten-day-old seedlings of WT, *OsLG3-OE* and RNAi plants and Digital Gene Expression (DGE) analysis was performed. Compared to WT, 223 transcripts were found to have a greater than 2-fold change in abundance (*P* < 0.05, FDR < 0.05) (**Figure 7a**). 159 genes were up-regulated in OE plants and down-regulated in RNAi plants, while 64 genes were down-regulated in OE plants and up-regulated in RNAi plants, respectively (**Figure 7b**). The expression of several of these genes was tested independent by qRT-PCR to validate the DGE results. Of the genes checked, six out of eight in OE plants (OE vs WT > 2) and five out of eight in RNAi plants (RNAi vs WT < 0.5) showed expression patterns consistent with the DGE results (**Additional file 6: Figure S6**). Gene Ontology (GO) analysis showed that the 218 differentially expressed genes affected by the *OsLG3* overexpression and suppression were significantly enriched in three GO terms (hyper geometric test, *P* <0.01, FDR<0.05), including response to stress (GO: 0006950), response to stimulus (GO: 0050896), and response to abiotic stimulus (GO: 0009628) (**Figure 7c; Additional file 3: Figure S3**). These results are consistent with our proposed role for *OsLG3* in the regulation of drought stress tolerance.

**Figure 7.**
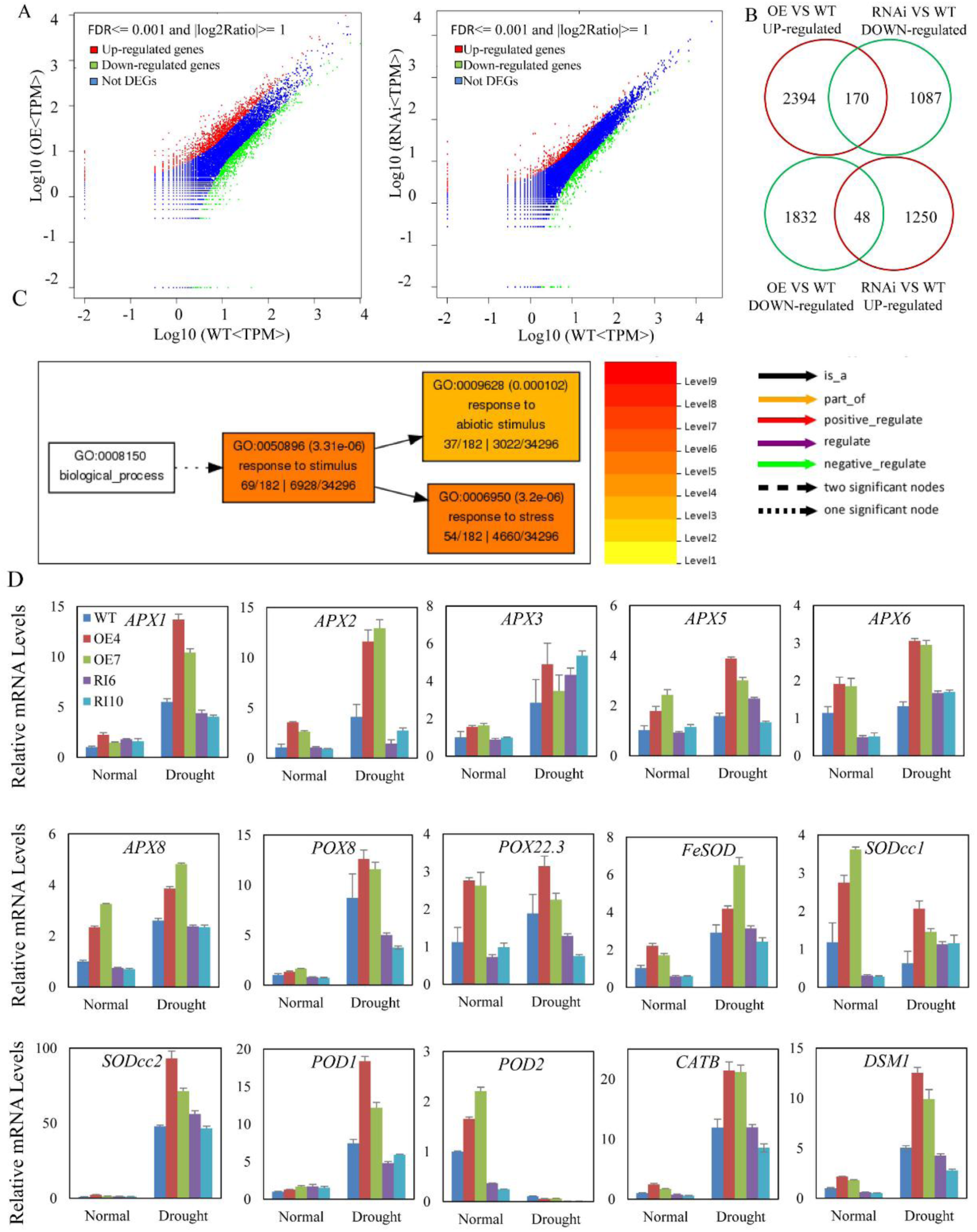
DGE analysis shows that altering *OsLG3* expression in transgenic plants affects transcription of stress response genes. **(a)** Scatterplots comparing the transcriptome of *OsLG3*-OE and -RNAi with WT. The red and green dots indicate transcripts from *OsLG3*-OE or -RNAi which have signal ratios compared to WT of greater than 2 and less than 0.5, respectively. **(b)** Venn diagram showing the numbers of up-regulated and down-regulated genes affected by the overexpression and suppression of *OsLG3*. **(c)** Significantly enriched GO terms show representative biological processes of up-regulated and down-regulated genes identified in *OsLG3*-OE and RNAi plants, respectively. **(d)** Transcript levels of genes related to ROS scavenging of WT, *OsLG3*-OE and RNAi plants under normal or drought stress conditions (withholding water for 5 days). Values are means ± SE (n = 3).

Interestingly, DGE analysis showed that ten ROS-scavenging related genes (*APX1*, *APX2*, *APX4*, *APX6*, *APX8*, *CATB*, *POD1*, *POD2*, *SODcc1*, *FeSOD*) were up-regulated in OE line and down-regulated in RNAi line. This result is consistent for a role of *OsLG3* in the control of ROS homeostasis. To confirm this regulation, we analyzed the expression levels of 15 genes by qRT-PCR (nine genes identified in the DGE analysis and six other ROS-related genes) in WT, *OsLG3*-OE and -RNAi plants under well-watered and drought conditions. 13 of the 15 tested genes were significantly up-regulated under drought stress conditions, and the expression level of 13 genes (except *APX3* and *POD2*) were significantly higher in OE plants than in WT and RNAi plants under drought stress conditions (**Figure 7d**). Conversely, the expression of *APX1*, *APX2*, *POX8*, *POX22.3* and *POD1* was significantly lower in the OsLG3-RNAi plants compared to WT under drought stress conditions, while the remaining nine genes did not show a significant difference in expression between WT and RNAi line (**Figure 7d**).

### *OsLG3* participates in H_2_O_2_ homeostasis

The potential role of *OsLG3* in oxidative stress tolerance was examined further by using two oxidative stress inducers, methyl viologen (MV) [56] and H_2_O_2_. Germinated WT and *OsLG3*-OE and RNAi plants were sown on ½ MS medium and ½ MS medium containing 2 μM MV. Application of MV dramatically repressed seedling growth in all plants, but the *OsLG3*-OE lines exhibited less growth inhibition compared to WT, while the RNAi plants showed more severe growth inhibition than WT (**Figure 8a, b and c**). Two-week-old seedlings were treated with 1 mM H_2_O_2_ or 3 μM MV for 24 hours, followed by 3, 3’-diaminobenzidine (DAB) staining to show the presence of H_2_O_2_ and nitroblue tetrazolium (NBT) staining to show the presence of superoxide anion. Under control conditions, WT or transgenic plants showed similar basal levels of H_2_O_2_ and superoxide, but DAB staining and NBT staining were much stronger in the WT than in the *OsLG3*-OE plants under H_2_O_2_ and MV stress treatment (**Figure 8d**). These results indicated that overexpression of *OsLG3* in rice can enhance tolerance to oxidative stress.

**Figure 8.**
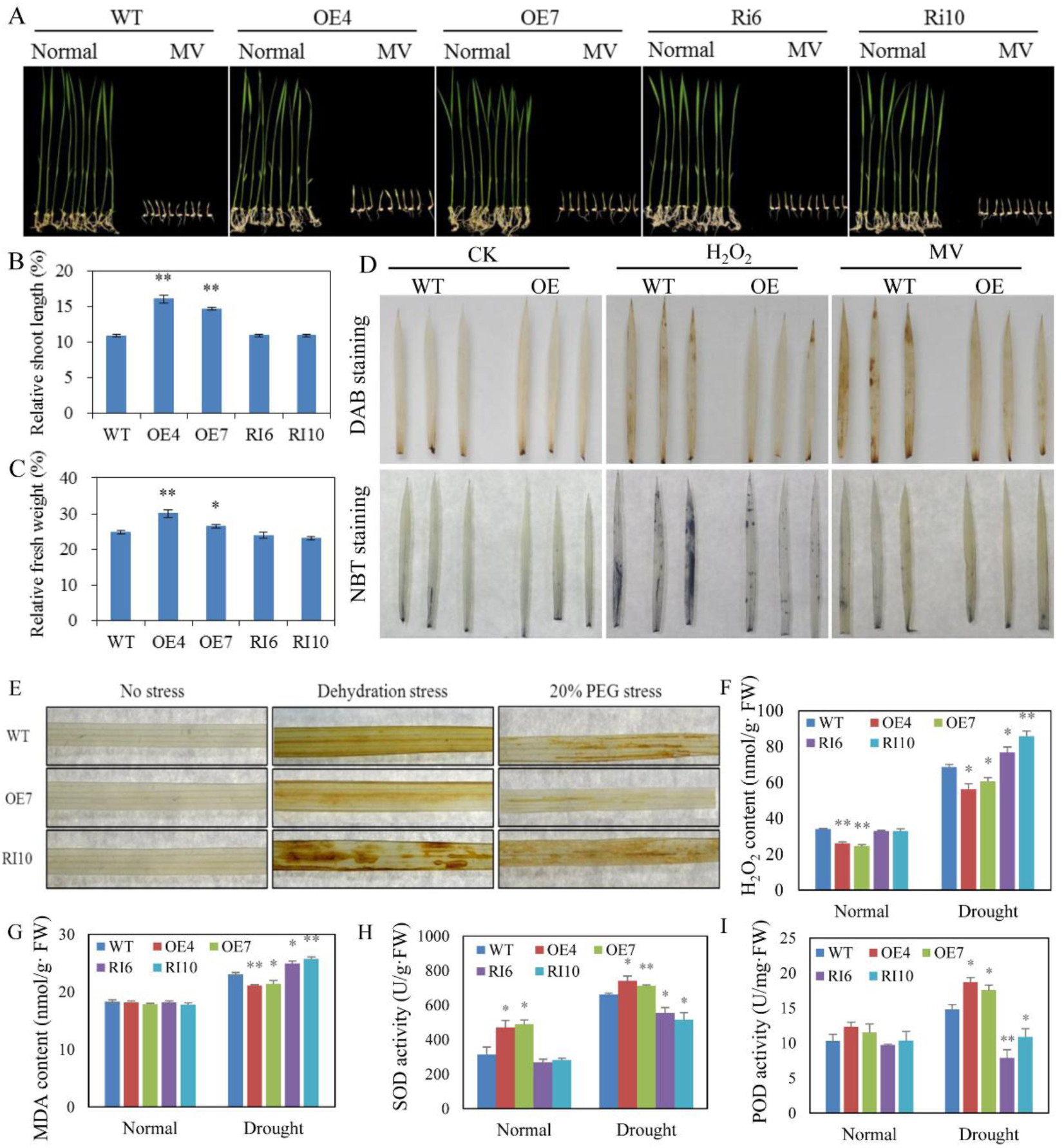
*OsLG3* is involved in oxidative stress response. **(a)** Enhanced tolerance of *OsLG3*-OE plants and enhanced sensitivity of *OsLG3*-RNAi plants to oxidative stress caused by methyl viologen (MV). Relative shoot length **(b)** and relative fresh weight **(c)** measurements of WT, *OsLG3*-OE and *OsLG3*-RNAi seedlings under oxidative stress treatments. Data are the mean ±SE (n=10). * and ** indicate significant differences at *P* < 0.05 and *P* < 0.01, respectively. **(d)** DAB and NBT staining of leaves for H_2_O_2_ in WT, *OsLG3-OE* and RNAi seedlings under oxidative stress treatments caused by H_2_O_2_ (100 mM) and MV (30 μM) stress treatment. **(e)** DAB staining of leaves for H_2_O_2_ from WT, *OsLG3*-OE and RNAi seedlings under normal conditions and stress treatment (three-week-old seedlings were treated with dehydration for 6 hours and 20% PEG6000 for 24 hours). **(f)** H_2_O_2_ content in leaves from WT, *OsLG3*-OE and RNAi seedlings under normal conditions and slight drought stress treatment (withholding water for 5 days). Relative MDA content **(g)**, SOD activity **(h)** and POD activity **(i)** in leaves from WT, *OsLG3*-OE and RNAi seedlings under normal and drought conditions (withholding water for 5 days). Data are the mean ± SE (n=3). * and ** indicate significant differences at *P* < 0. 05 and *P* < 0.01, respectively.

Based on the above results, it is hypothesized that *OsLG3* may be involved in the regulation of ROS homeostasis, leading to enhanced adaption to drought. To confirm this, we stressed the WT, *OsLG3*-OE and -RNAi plants with dehydration and 20% PEG, followed by DAB staining to detect H_2_O_2_ accumulation. Under well-watered conditions, there was low H_2_O_2_ accumulation level in WT, *OsLG3*-OE and RNAi plants (**Figure 8e**). However, *OsLG3*-OE leaves showed significantly less accumulation of H_2_O_2_ than WT plants under drought stress conditions, whereas RNAi lines shows more H_2_O_2_ than WT under drought stress. H_2_O_2_ accumulation under normal and drought stress conditions was also quantified, supporting the DAB staining results. Less H_2_O_2_ was detected in the leaves of *OsLG3*-OE plants than WT and *OsLG3*-RNAi plants under drought stress conditions (**Figure 8f**). The accumulation of ROS may lead to oxidative damage, as indicated by membrane lipid peroxidation. Monodehydroascorbate (MDA) is a biomarker for membrane lipid peroxidation [57-59]. MDA production under normal growth conditions was similar in WT and all transgenic plants, whereas under drought stress, MDA production was significantly lower in the *OsLG3*-OE compared with WT and OsLG3-RNAi plants (**Figure 8g**). These results demonstrate that overexpression of *OsLG3* can reduce the damage to membranes caused by drought stress.

The effect of *OsLG3*-OE on oxidative stress tolerance implies *OsLG3* may influence ROS homeostasis. It is well known that superoxide dismutase (SOD), and peroxidase (POD) are the major ROS-scavenging enzymes in plants under stressed conditions [58]. Four-week-old seedlings were treated with drought stress for 5 days and the activity of SOD and POD was determined. Results suggested that under normal growth conditions, *OsLG3*-OE lines have significant higher SOD than WT and RNAi plants (**Figure 8h**), while the activity of POD did not appear to be significantly affected in *OsLG3*-OE or RNAi plants (**Figure 8i**). Under drought stress conditions, the activities of POD and SOD were both significantly enhanced in OE plants and significantly reduced in RNAi plants compared to WT (**Figure 8h, i**). These results imply that the function of *OsLG3* in drought tolerance may be associated with an enhanced antioxidant response to counteract oxidative stress under drought.

### The elite *OsLG3* allele can be used for improving *japonica* rice drought tolerance

To analyze the relationship between *OsLG3* haplotypes and drought tolerance, we investigated the phylogenetic relationship of 1058 deep-sequenced (depths**≈**14.9x) rice accessions, including 251 UR, 415 LR and 392 wild rice varieties originating from a wide geographic range (**Additional file 10: Data Set 1**). Forty-five SNPs were identified by minor allele frequency (MAF) > 0.05 and missing rates ☐**≤**☐50%. Phylogenetic analysis based on these variations showed that there is a clear differentiation between *japonica*-UR and *japonica*-LR rice (**Figure 9a**). To get further insight into the phylogenetic relationship, ten elite *japonica*_UR varieties (IRAT109, IAC150/76, IRAT266, Guangkexiangnuo, Shanjiugu, Taitung_upland328, Jaeraeryukdo, Riku aikoku, Padi darawal, Malandi 2) were chosen as an UR pool based on their strong drought resistance, and ten typical *japonica*-LR varieties (Nipponbare, Yuefu, Guichao2, Koshihikari, Zhonghua11, Early_chongjin, Xiushui115, IR24, Ningjing3, Xiushui114) were chosen as a LR pool. Seven SNPs (SNP_4352414, SNP_4352886, SNP_4352960, SNP_4352792, SNP_4352797, SNP_4353103 and SNP_4353347) that differed between the upland rice pool and lowland rice pool were identified. These seven SNPS showed a clear phylogenetic distinction between *japonica*-UR and *japonica*-LR (**Figure 9b**). On the basis of these seven SNPs, we could divide the sequences of the 1058 cultivated varieties into nine haplotypes of which there were four main variants, Hap1, Hap2, Hap3, and Hap4 (**Figure 9c, d**). *OsLG3*^Nipponbare^ is representative of Hap1, which is mainly composed of *japonica*-LR rice, whereas *OsLG3*^IRAT109^ belongs to Hap2, which is mainly composed of *japonica*-UR rice and is the second largest group. Hap3 is mainly composed of *indica* rice, and Hap4 is mainly composed of wild rice. The results indicated that *OsLG3* has clear differentiation in *japonica* rice, which can be divided into *japonica*-UR and *japonica*-LR rice, while no clear division in *indica* rice. Therefore, we designated Hap1 and Hap2 as the sensitive and tolerant alleles, respectively, of *OsLG3* in *japonica* rice.

**Figure 9.**
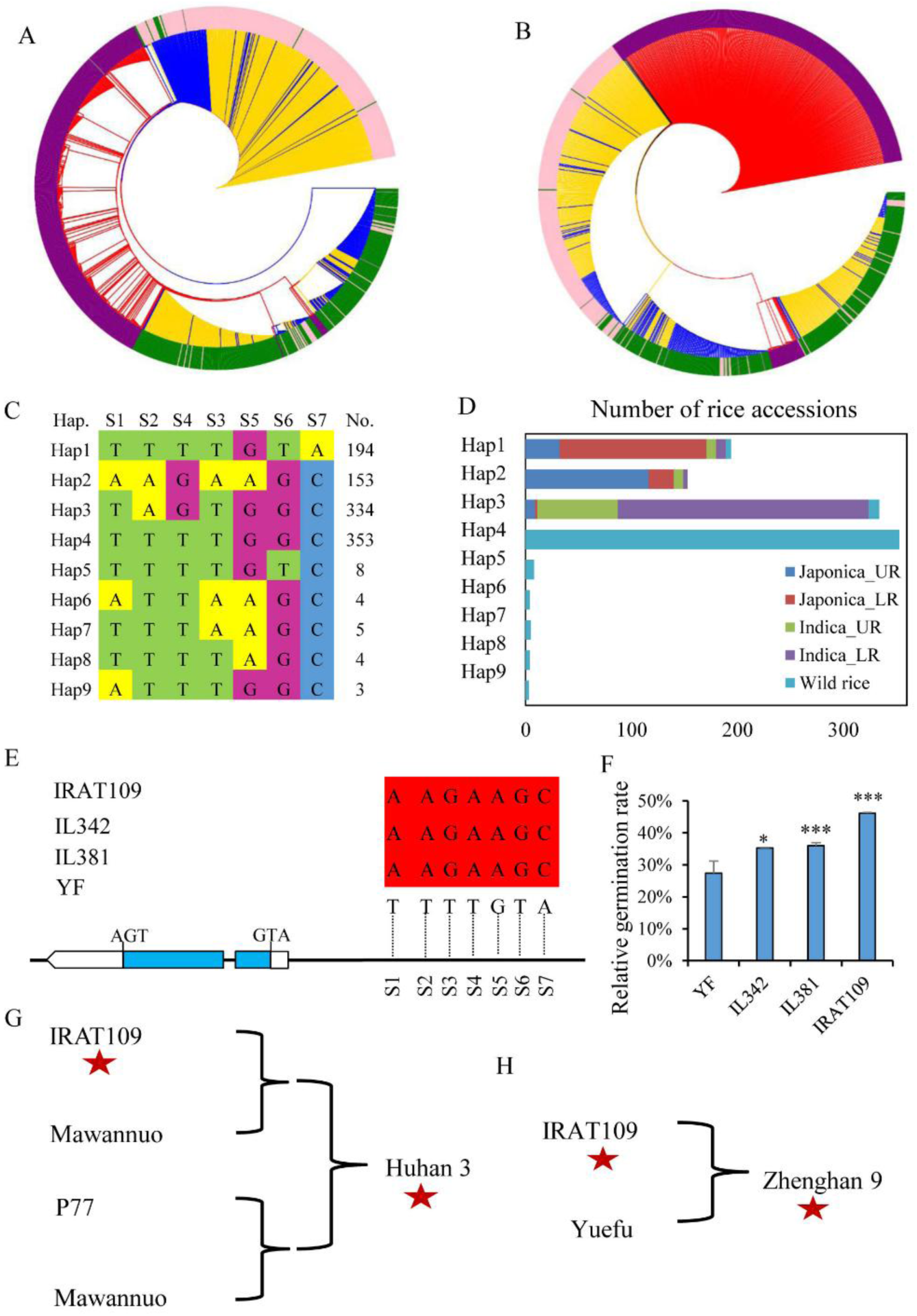
The favorable allele of *OsLG3* improves drought tolerance in rice. **(a)** Phylogenetic analysis of the *OsLG3* gene based on 45 SNPs in 1058 accessions. **(b)** Phylogenetic tree of the *OsLG3* in 1058 varieties was constructed based on seven SNPs (SNP_4352414, SNP_4352792, SNP_4352797, SNP_4352886, SNP_4352960, SNP_4353103 and SNP_4353347). Different colors reflect the different subgroups. The pink and green strips represent *indica* and *japonica*, respectively. Both purple strip and red lines represent wild rice accessions. The blue and gold lines indicate upland rice and lowland rice, respectively. **(c, d)** Haplotype analysis of *OsLG3*. Natural variation in *OsLG3* among 1058 rice accessions of a worldwide rice collection. S1-7 denote SNP_4352414, SNP_4352792, SNP_4352797, SNP_4352886, SNP_4352960, SNP_4353103 and SNP_4353347, respectively. Hap., haplotype; No., number of cultivated varieties; *Japonica*_UR, upland rice in *japonica; Japonica*_LR, lowland rice in *japonica; Indica*_UR, upland rice in *indica; Indica*_LR, lowland rice in *indica*. **(e)** Haplotypes of *OsLG3* in IL342, IL381, IRAT109 and Yuefu rice genotypes. ATG is the site of the start codon. SNP4 is incomplete LD with other six polymorphisms in the three drought tolerant lines. These polymorphisms are shaded in red. **(f)** Relative germination rate of IL342, IL381, IRAT109 and Yuefu rice genotypes. Data represent the mean of triplicates (\**P* < 0.05, ** *P* < 0.01). **(g, h)** The pedigree of selected rice varieties. **(g)** Huhan 3 and **(h)** Zhenghan 9. The red star refers the beneficial allele of *OsLG3*.

To confirm the genetic effect of different alleles of *OsLG3* on rice drought tolerance, two introgression lines (IL342 and IL381) were selected from a cross between IRAT109 (donor parent) and Yuefu (receptor parent). The *OsLG3* allele in these two lines shares sequence similarity with IRAT109, with seven matching SNPs in the promoter (including the three significant loci describe in candidate gene association analysis section), whereas Yuefu carries the same *OsLG3* allele as Nipponbare (**Figure 9e**). Thus, the *OsLG3* allele found in IRAT109, IL342 and IL381 was considered to be the drought-tolerant allele and the allele in Yuefu the drought-sensitive allele, respectively. When germinated on 15% PEG conditions, the RGR of Yuefu was 27.0% compared to germination on water, while the RGR of IL342, IL381 and IRAT109 was 35%, 36% and 46%, respectively (**Figure 9f**). These results support the hypothesis that the nucleotide polymorphisms in *OsLG3* contribute to enhanced rice drought tolerance.

According to the pedigree records, Huhan3, one of green super rice accessions, is an elite upland rice that is widely grown in Hubei province of China because of its high yield and outstanding ability to conserve water. One of its parents is IRAT109. A re-sequencing study showed that Huhan3 carries the allele of *OsLG3* deriving from IRAT109 (**Figure 9g**). Moreover, Zhenghan9, another elite upland rice, is derived from the cross of IRAT109 and Yuefu. Zhenghan9 retained the favorable allele of *OsLG3* and an allele for an unidentified grain quality-related gene derived from Yuefu (**Figure 9h**). These observations illustrate the successful outcome of combining the *OsLG3^IRAT109^* allele and those alleles for other unidentified grain quality-related genes, for improving drought tolerance and grain appearance quality, which had been employed by breeders.

## Discussion

### Natural variation in the promoter of *OsLG3* is associated with drought tolerance in rice

Despite many studies investigating the transcriptional response to drought stress [5, 60-62], how natural sequence variation is associated with phenotypic variations in drought tolerance remains largely unknown. Because of the polygenic inheritance, low heritability, and strong genotype-by-environment interactions of drought resistance-related traits, there are few reports on the cloning and identification of drought resistant genes by association analysis and positive mutant screening methods [48, 49, 63]. In this study, using candidate gene association analysis, we detected three significant SNPs (*P* < 1.0 × 10^-3^) (SNP_4352414, SNP_4352886, SNP_4352960) located within the promoter region of *OsLG3* were significantly associated with the RGR trait (**Figure 1c**). From our previous work, transient assays showed that four out of seven SNPs are conserved and play a significant role in regulating *OsLG3* expression [51]. In this study, transgenic experiments further support the hypothesis that increased expression of *OsLG3* enhances rice drought stress tolerance.

To analyze the relationship between *OsLG3* haplotypes and drought tolerance, we investigated the phylogenetic relationship of 1058 deep-sequenced rice accessions. Based on the seven major SNPs, we divide the sequences of the 1058 cultivated varieties into nine haplotypes of which there were four main variants, Hap1 (mainly composed of *japonica*-LR rice), Hap2 mainly composed of *japonica*-UR rice, Hap3 (mainly composed of *indica* rice), and Hap4 (mainly composed of wild rice). The variations of *OsLG3* in these accessions showed that there is a clear differentiation between *japonica*-UR and *japonica*-LR rice, while no clear division in *indica* and wild rice (**Figure 9**). These results implicated the Hap2 variant of *OsLG3* may improve drought tolerance of cultivated *japonica* rice. Re-analysis of two introgression lines (IL342 and IL381) derived from a cross between IRAT109 (donor parent) and Yuefu (receptor parent) indicated that selection of Hap2 was effective in the improvement of drought tolerance (**Figure 9e**). Interestingly, our previous work indicated that *OsLG3* acts as an important positive regulator of grain length and could improve rice yield [51]. In fact, the pedigree records in upland rice breeding showed that the tolerant allele of *OsLG3* had been incorporated into elite varieties via breeding (**Figure 9g**). We propose that pyramiding the beneficial allele of *OsLG3* with other yield- and quality-related genes could contribute to the breeding of elite rice varieties because of its pleiotropic effects on traits.

### *OsLG3* was cloned from upland rice and plays a positive role in rice drought tolerance

UR has evolved enhanced drought-resistance compared to LR, derived from natural and artificial selection over time under drought conditions. UR performs better under drought conditions, with stronger water-retention ability, larger root volumes and higher biomass production [64, 65]. Therefore, UR is highly suitable to be taken up as research material regarding mechanisms underlying drought resistance. From the previous work of expression profiles from typical LR (Nipponbare and Yuefu, drought-sensitive *japonica* rice) and UR varieties (IRAT109 and Haogelao, drought-resistant *japonica* rice) under water deficit stress conditions using cDNA microarray [52], we found the transcription of *OsLG3* can be induced to a greater extent in the UR varieties compared to the LR varieties during drought stress. When IRAT109 and Nipponbare were subjected to soil drought stress, expression of *OsLG3* in Nipponbare was not induced, whereas in IRAT109 *OsLG3* shows significant induction under both slight drought (SLD) and moderate drought (MOD) conditions (**Figure 1a**). Using candidate gene association analysis, we found that nucleotide polymorphisms in the promoter region of *OsLG3* are associated with different levels of water deficit tolerance among rice varieties at the germination stage (**Figure 1b**). All these results indicated that *OsLG3* might play a role in the observed drought stress response of upland rice.

To assess the effect of *OsLG3* on water deficit stress responses, we tested the growth response of *OsLG3*-OE and -RNAi plants under simulated drought stresses. *OsLG3*-OE lines showed higher survival under dehydration caused by 20% PEG6000 and soil drought stress caused by stopping watering (**Figure 5**). Plants with reduced *OsLG3* expression showed reduced survival, suggesting that this gene might be involved in response to drought stress. A slower rate of water loss was observed in the leaves of *OsLG3*-OE compared to WT and RNAi plants (**Additional file 5: Figure S5a**), suggesting that *OsLG3* plays an important role in reducing water evaporation. Collectively, these results demonstrate that *OsLG3* is a positive regulator of drought stress response in rice. *OsLG3*-OE plants also showed enhanced growth compared to WT and RNAi lines under mannitol and NaCl treatments, which suggests that *OsLG3* may be involved in osmotic and high salinity stress tolerance through cross-talk between water deficit, osmotic and high salinity stress response pathways.

### Potential mechanisms of *OsLG3* in drought stress tolerance

We found that the expression levels of a set of stress-related genes like *OsLEA3*, *OsAP37*, *SNAC1* are increased in *OsLG3*-OE lines compared to WT, and decreased in *OsLG3*-RNAi lines before and after drought treatment (**Additional file 5: Figure S5b**). This was confirmed by DGE analysis (**Additional file 9: Table S3**). We investigated if the altered response to drought was associated with global changes in the expression of stress-related genes before stress application. Analyses indicated that many stress-related genes were up-regulated in the OE line and down-regulated in the RNAi line (**Figure 7c**), which is consistent with the observed phenotypes of *OsLG3*-OE and -RNAi lines. Interestingly, the expression level of some ROS-scavenging related genes showed increased expression levels in OE and decreased abundance in RNAi lines too. For instance, the ascorbate peroxidase gene, *OsAPX1* plays a positive role in chilling tolerance by enhancing H_2_O_2_-scavenging [66]. *DSM1* mediates drought resistance through ROS scavenging in rice [12]. *OsCATB* prevent the excessive accumulation of H_2_O_2_ under water stress [67]. Thus, these DGE analyses are consistent with our hypothesis that the function of *OsLG3* in abiotic stress tolerance occurs through the transcriptional regulation of stress-related and ROS scavenging-related genes.

Previous studies have demonstrated that plant response to abiotic stresses is mediated through the regulation of ROS metabolism [9, 10, 12, 68, 69]. For example, overexpression of a NAC gene, *SNAC3*, increased drought and heat tolerance by modulating ROS homeostasis through regulating the expression of genes for ROS-scavenging and production enzymes [9]. Overexpression of an ERF gene, *SERF1*, improved salinity tolerance mainly due to the regulation of ROS-dependent signaling during the initial phase of salt stress in rice [10]. The present data revealed overexpression of *OsLG3* led to the up-regulation of many ROS scavenge-associated genes, while suppression of *OsLG3* caused the down-regulation of these genes (**Figure 7d**). Furthermore, the H_2_O_2_ and MDA content which accumulated in the leaves of *OsLG3*-OE plants was significantly lower than that in the WT and *OsLG3*-RNAi plants (**Figure 8c, d**), suggesting that the improved drought tolerance of *OsLG3*-OE plants may be due to the efficient scavenging of ROS. Relevant to this, *OsLG3*-OE seedlings exhibited better growth under oxidative stress caused by application of MV. In contrast, RNAi plants showed enhanced sensitivity to oxidative stress. Therefore, it is proposed that the function of *OsLG3* in drought tolerance is associated with the enhancing activity of ROS-scavenging.

To scavenge or detoxify excess stress-induced ROS, plants have developed a complex antioxidant system comprising of the non-enzymatic as well as enzymatic antioxidants [7, 70]. The POD and SOD activities were found to be higher in the *OsLG3*-OE lines as compared to WT under drought stress, and *OsLG3*-RNAi lines showed the reverse results (**Figure 8h**). These data suggest that *OsLG3*-OE plants have enhanced activity of ROS-scavenging enzymes which significantly contributes to the reduction of ROS accumulation, and thereby improved drought stress tolerance.

In conclusion, here we present evidence that *OsLG3* is induced by water deficit stresses and its induction is greater in UR IRAT109 compared to LR Nipponbare under drought. Nucleotide polymorphisms in the promoter region of *OsLG3* are associated with water deficit tolerance in germinating rice. Transgenic plants overexpressing *OsLG3* showed improved growth under drought stress, probably via inducing ROS scavenging by controlling downstream ROS-related genes. Phylogenetic analysis indicated that the tolerant allele of *OsLG3*, identified in drought-tolerant *japonica* rice varieties, could be introduced to rice to improve the drought tolerance. The pedigree records in upland rice breeding showed that the tolerant allele of *OsLG3* had been incorporated into elite varieties via breeding. Importantly, the tolerant allele of *OsLG3* is a promising genetic resource for the development of drought-tolerant and high yield rice varieties using traditional breeding approaches or genetic engineering.

## Methods

### Plant materials and stress treatments

Rice varieties IRAT109 and Nipponbare were used for quantitative real-time PCR (qRT-PCR) analysis of *OsLG3* transcript levels under various stresses and hormone treatments, and Nipponbare was used for all transgenic experiments. For qRT-PCR analysis the expression level of *OsLG3* under simulated drought conditions, the seeds of Nipponbare and IRAT109 were sown in flower pots (140 mm diameter×160 mm deep) with well-mixed soil (forest soil: vermiculite in a ratio of 1:1), and grown in the greenhouse under well water conditions at 28°C/ 26°C and 12 hours light/12 hours dark photoperiod. Three-week old plants were subjected to drought treatment with stop watering for 0, 5, 6 and 7 days. From these, four stress treatment levels were categorized as no stress (NS), slight drought (SLD), moderate drought (MOD) and severe drought (SED), respectively. For qRT-PCR analysis the expression level of *OsLG3* under various abiotic stresses and phytohormone treatments, three-week-old IRAT109 seedlings grown in Hoagland solution (PPFD of 400 μmol/m^2^s^2^, 12 hours light (28°C)/12 hours dark (26°C)) were subjected to different treatments with 20% (w/v) polyethylene glycol (PEG) 6000, NaCl (200 mM), cold (4°C), hydrogen peroxide (H_2_O_2_,1 mM), abscisic acid (ABA,100 μM), ethylene (ETH,100 μM), GA (100 μM) and dehydration by exposure in air. Leaf tissue was harvested at 0, 1, 2, 4, 6, 9, 12 and 24 hours after PEG, NaCl, cold, H_2_O_2_, ABA, ETH, and GA treatments, and at 0, 1, 2, 3, 4, 5, 6, 7 and 8 hours after dehydration treatment.

The seeds of WT, *OsLG3*-OE lines (OE4, OE7) and *OsLG3*-RNAi lines (RI6, RI10) were germinated in ½ MS medium for various stress evaluations. For dehydration treatment, uniformly germinated seeds were transplanted into bottom removed 96-well PCR plates, and grown hydroponically using Hoagland solution at 28°C/26°C (day/night) with a 12 hours photoperiod. Three-week-old plants were treated with 20% (w/v) PEG6000 solution for about three days, and recovered with water for ten days. Each stress test was repeated three times. For drought stress treatment, three-day-old WT and transgenic seedlings were transplanted to well-mixed soil (forest soil: vermiculite in a ratio of 1:1), and grown for four weeks under a normal watering regime. Drought stress treatment was then applied by stopping irrigation for about seven days. After all leaves had completely rolled, watering was resumed for 10 days. The survival ratio of each line was calculated as the number of surviving plants over the number of treated plants in each flowerpot. To evaluate the tolerance of rice seedlings to osmotic, high salinity and oxidative stress treatment, three-day-old WT, *OsLG3*-OE and *OsLG3*-RNAi seedlings (10 plants per replicate, three replicates) were transplanted to ½ MS medium (mock treatment) or ½ MS medium containing 200 mM mannitol, 150 mM NaCl or 2 μM MV respectively. Plants were grown for seven days under 12 hours light (28°C)/12 hours dark (26°C) photoperiod conditions and then the shoot height and fresh weight of all plants was measured.

### Gene expression analysis

Total RNA was extracted using RNAiso Plus (TaKaRa) according to the manufacturer’s instructions. 4 μg of the DNase-treated RNA were reverse transcribed by using M-MLV reverse transcriptase (TaKaRa). qRT-PCR was performed in an Applied Biosystems 7500 Real Time PCR system (ABI, USA) using SYBR Premix Ex TaqTM II (TaKaRa) according to the protocol previously described [71, 72]. The gene-specific primers used for qRT-PCR are listed in **Supplemental Table S2**. Rice *Actin1* gene was used as the internal reference gene for data normalization [73].

### *OsLG3*-gene association analysis of rice drought tolerance among 173 rice genotypes

173 cultivated varieties from the mini core collection of Chinese cultivated rice (Zhang et al. 2010), including 130 *indica* and 43 *japonica* rice varieties (**Supplemental Table S1**) were selected for the candidate gene association mapping. To perform the analysis, we obtained phenotypic data of relative germination rates (RGR, ratio of germination rates under stress conditions to germination rates under water conditions) from growth in water and 15% PEG6000 treatment. Briefly, 50 seeds of each line were placed in petri dishes (90 mm diam) lined with filter paper. 10 ml water or 15% PEG6000 was added to each petri dish as mock treatment or to induce osmotic stress, respectively. All petri dishes were placed in a 28°C greenhouse and the germination rates were assessed after five days. Genotype data for each line was acquired from ‘The 3,000 rice genomes project’ [74]. The SNP data were filtered out. After filtering, a total of 97 SNPs remained in a 5-kb region surrounding the *OsLG3* gene. Association analysis using a general linear model with the population structure (Q matrix) method was conducted using TASSEL 5.2.28 software. The Q matrix was estimated from the genomic data to control for population structure.

### Plasmid construction and rice transformation

To generate the overexpression construct, the full-length cDNA of *OsLG3* was amplified from the first-strand cDNA of IRAT109 with specific primers (**Supplemental Table S2**) using PrimeSTAR^®^ HS DNA Polymerase (TaKaRa), digested with *Kpn*I and *Pac*I, and cloned into binary vector pMDC32 [75]. For construction of RNA interference (RNAi) plasmid, a 415 bp fragment with low similarity to other rice genes located at the 5’ end of *OsLG3* was amplified by PCR with specific primer, digested with *Kpn*I and *Bam*HI and then *Spe*I and *Sac*I sites, and cloned into the pTCK303 vector as described previously [76].

### Subcellular localization

For subcellular localization analysis, the full-length open reading frame (ORF) of *OsLG3* without terminal codon was amplified, and the amplified fragments were digested with *Kpn*I and *Pac*I, and then cloned in to pMDC83 vector, fused with the GFP reporter gene driven by *CaMV 35S* promoter. The fusion construct (35S:OsLG3-GFP) and control (35S:GFP) were transformed into *Agrobacterium tumefaciens* strains (EH105), and then infiltrated to five-week-old *Nicotiana benthamiana* leaves [77]. The fluorescence signal was examined through a confocal laser scanning microscope OLYMPUS FV1000 with excitation at 488 nm and emission at 525 nm.

### Transactivation and dimerization assay

For transactivation assay, The full length coding region and truncated fragments of *OsLG3* generated by PCR amplification were fused in frame to the GAL4 DNA binding domain in the vector of pGBKT7 (Invitrogen). The plasmids of FL (the full length coding region of OsLG3, amino acids 1-334), dC1 (amino acids 1-218), dC2 (amino acids 1-167), dC3 (amino acids 1-106), dC4 (amino acids 107-334), dC5 (amino acids 168-334) and dC6 (amino acids 213-334) were constructed. These constructs were introduced into yeast strain AH109 by LiAc-mediated yeast transformations, and screened on the selective medium plates without Tryptophan (SD/-Trp). The PCR-verified transformants were transferred to SD medium without Tryptophan/Histidine/Adenine (SD/-Trp/-His/-Ade) for 3 days. The β-galactosidase activity was performed according to the *in vivo* agar plate assay (X-α-gal in medium).

For dimerization assay, AD-OsLG3 with full length coding region of *OsLG3* fused in frame to the GAL4 DNA activating domain in the vector of pGADT7 (Invitrogen) was constructed. The constructs AD-OsLG3 and BD-dC2 were co-transformed into yeast strain AH109. AD and BD empty, AD-OsLG3 and BD empty were also co-transformed as negative controls. All transformed cells were screened on the selective medium plates without Tryptophan and Leucine (SD/-Trp-Leu), The PCR-verified transformants were transferred to SD medium with X-α-gal and without Tryptophan/Histidine/Adenine/Leucine (SD/-Trp-Leu-Ade-His/X-α-gal) for 3 days.

### Physiological measurements

The rate of water loss under dehydration condition was measured as described previously [72]. Ten plants of each line were used in each replicate, and three replicates were made for each line. Histochemical assays for ROS accumulation were determined according to the previously described method [68, 78]. Briefly, the qualitative detection of H_2_O_2_ accumulation was detected by DAB staining. Excised leaves were treated with DAB staining solution (1mg/mL DAB pH 3.8) at 28°C for 12 hours in the dark. After staining, the leaves were decolorized with Acetic acid/Ethanol (1:3) for 60 min, and rehydrated in 70% (v/v) alcohol for 24 hours at 28°C. Each experiment was repeated on at least 10 different plants, and representative images were shown. Superoxide anion radical accumulation was detected by NBT staining as described previously. The leaf samples were excised and immediately placed in 50 mM sodium phosphate buffer (pH 7.5) containing 6 mM NBT, at 28°C for 8 h in the dark. The quantitative measurement of H_2_O_2_ concentrations was performed with an Amplex Red Hydrogen / Peroxidase Assay Kit (Molecular Probes) (Invitrogen, http://www.invitrogen.com) as described by the manufacturer’s instructions. Briefly, leaf samples from both the well water and drought stressed (without water for 5 days) plants were ground in liquid nitrogen, and 100 mg of ground frozen tissue from each sample was placed in a 2 mL Eppendorf tube and kept frozen. 1 mL precooled sodium phosphate buffer (20 mM, pH 6.5) was immediately added into the tube and mixed. After centrifugation (10,000 g, 4°C, 10 min) the supernatant was used for the assay. Measurements were performed using a 96-well microplate reader (PowerWave XS2, BioTek) at an absorbance of 560 nm. MDA content was measured as described previously [72].

The activity of antioxidant enzymes including superoxide dismutase (SOD) and peroxidase (POD) were measured following the protocols described previously [71]. The units of the antioxidant enzyme activities were defined as follows: One unit of the SOD activity was defined as the quantity of enzyme required to cause 50% inhibition of the photochemical reduction of NBT per minute at 560 nm. One unit of POD activity was defined as the amount enzyme required to cause a 0.01 absorbance increase per minute at 470 nm.

### Digital gene expression (DGE) profiling and gene ontology (GO) analysis

WT Nipponbare and transgenic lines OE7 and RI10 were used for digital gene expression (DGE) profiling analysis. Ten-day-old seedlings grown on ½ MS medium were harvested for total RNA extraction as described in qRT-PCR section. DGE was performed at the Beijing Genomics Institute (http://www.genomics.cn) using Illumina Hiseq2000 sequencing technology. Transcripts with significant differential expression between *OsLG3*-OE and WT or between *OsLG3*-RNAi and WT plants were identified as those with *P* < 0.05 using a False Discovery Rate (FDR) of < 0.05 and a fold change cutoff of >2. Gene Ontology (GO) analysis was performed using agriGO [79] (http://bioinfo.cau.edu.cn/agriGO/). Representative differentially expressed genes were confirmed by qRT-PCR. The primers are listed in **Supplemental Table S2**.

### Sequence analysis and phylogenetic analysis

The phylogenetic tree was analyzed by MEGA6 software based on neighbor-joining method and bootstrap analysis (1,000 replicates). The EvolView online tool [80] was used for visualizing the phylogenetic tree. Multiple sequence alignment was performed with ClustalW. Gene annotation was conducted in RGAP (http://rice.plantbiology.msu.edu/). All SNP data were obtained from the Rice functional genomics and breeding database (RFGB, http://www.rmbreeding.cn/Index/).

### Accession numbers

Sequence data from this article can be found in the Rice Genome Annotation Project Database and Resource (RGAP) (http://rice.plantbiology.msu.edu) under following accession numbers: *OsLG3* (LOC_Os03g08470); *Actin1* (LOC_Os10g36650); *OsLEA3* (LOC_Os05g46480); *AP37* (LOC_Os01g58420); *SNAC1* (LOC_Os03g60080); *RAB21* (LOC_Os11g26790); *RAB16C* (LOC_Os11g26760); *OsbZIP23* (LOC_Os02g52780); *Apx1* (LOC_Os03g17690); *Apx2* (LOC_Os07g49400); *Apx3* (LOC_Os04g14680); *Apx5* (LOC_Os12g07830); *Apx6* (LOC_Os12g07820); *Apx8* (LOC_Os02g34810); *OsPox8.1* (LOC_Os07g48010); *Pox22.3* (LOC_Os07g48020); *FeSOD* (LOC_Os06g05110); *SODcc1* (LOC_Os03g22810); *SODcc2* (LOC_Os07g46990); *POD1* (LOC_Os01g22370); *POD2* (LOC_Os03g22010); *CATB* (LOC_Os06g51150); *DSM1* (LOC_Os02g50970); *RAB16D* (LOC_Os11g26750); *OsNCED4* (LOC_Os07g05940); *OsNCED3* (LOC_Os03g44380); *OsDhn1* (LOC_Os02g44870); *OsSRO1C* (LOC_Os03g12820); *OsMYB48* (LOC_Os01g74410); *OsITPK4* (LOC_Os02g26720); *OsLEA3-2* (LOC_Os03g20680).

## Author’s contributions

H. X. and J. Y designed and performed most of the research and wrote the article, X. W. and P. L. contributed to help with transgenic experiment and stress treatment, J.L., H. Z. and Y. G. contributed to design some experiments, Y. Z. contributed to help with haplotype analysis, Y. L and Z. Y. contributed to sample or reagents preparation, B. F., W. W., J. A. and Z. L. help to analyze the data and revise the manuscript, and Z.L conceived the research and assisted in writing the manuscript.

## Acknowledgments

We thank Amelia Henry (International Rice Research Institute), Steven Burgess (eLIFE) and Andrew Plackett (University of Cambridge) for critical reading and suggest revisions to this manuscript.

## Additional files

**Additional file 1: Figure S1.** Frequency distribution of relative germination rates in the mini core collection (MCC) population.

**Additional file 2: Figure S2.** Sequence analysis of OsLG3. **(a)** Schematic diagram of OsLG3 gene structure. Representation of OsLG3 protein domain structure showing the location of the AP2 domain and the DNA-binding sites (indicated by shaded triangles). **(b)** Multiple sequence alignment of OsLG3 with several previously reported stress-related ERF genes in rice with DNAMAN software. Accession numbers are as follows: OsERF71: *Oryza sativa*, LOC_Os06g09390; OsBIERF1: *Oryza sativa*, LOC_Os09g26420; Sub1A: *Oryza sativa*, DQ011598; OsEREBP1: *Oryza sativa*, LOC_Os02g54160; AtEBP: *Arabidopsis thaliana*, AT3G16770; CaPF1: *Capsicum annuum*, AAP72289; GmEREBP1: *Glycine max*, NP_001236578; JERF1: *Solanum lycopersicum*, NP_001234513. JERF3: *Solanum lycopersicum*, AAQ91334. Red line, black line and red box represent MCGGAIL/I motif, nuclear localization signal and conserved AP2 domain, respectively. **(c)** Sequence logos of the AP2 domain were produced based on the sequences presented in **(b)**. Height of letter (amino acid) at each position indicates degree of conservation.

**Additional file 3: Figure S3.** Overexpression of *OsLG3*. **(a)** *OsLG3* overexpression construct used for rice transformation. LB, left border; HPT, *Hygromycin phosphotransferase;* 35S, *cauliflower mosaic virus 35S* promoter; RB, right border. **(b)** The expression level of *OsLG3* in WT and OE lines analyzed by qRT-PCR. The lines used for further analysis were labeled in black star. **(c)** Phenotype of *OsLG3*-OE and WT plants under normal growth conditions.

**Additional file 4: Figure S4.** Suppression of *OsLG3* by RNAi. **(a)** *OsLG3*-RNAi construct used for rice transformation. **(b)** The expression level of *OsLG3* in WT and RNAi lines analyzed by qRT-PCR. The lines used for further analysis were labeled in black star. **(c)** Phenotype of WT and *OsLG3*-RNAi plants under normal growth conditions at seedling stage.

**Additional file 5: Figure S5.** Water loss from detached leaves and expression of drought stress-related genes in WT, *OsLG3*-OE and RNAi plants under normal and drought stress conditions. **(a)** Water loss from detached leaves of WT, *OsLG3*-OE and -RNAi plants at indicated time points. Data are the mean ± SE (n=3). * and ** indicate significant differences at *P* < 0.05 and *P*< 0.01, respectively. **(b)** Expression of drought stress responsive genes in WT, *OsLG3*-OE and RNAi under normal and drought stress conditions (without water for 5 days). Values are means ± SE (n=3).

**Additional file 6: Figure S6.** qRT-PCR validation of differentially expressed genes in *OsLG3*-OE **(a)** and *OsLG3*-RNAi transgenic plants **(b)** compared to WT Plants.

**Additional file 7: Table S1.** Information of 173 MCC varieties used for association analysis.

**Additional file 8: Table S2.** Primer sequences used in this study.

**Additional file 9: Table S3.** GO_biological_process analysis of the differentially expressed genes.

**Additional file 10: Data Set 1.** Information of 1058 *Oryza sativa* varieties and wild rice used for haplotype analysis.

**Additional file 11: Data Set 2.** Differentially expressed genes in *OsLG3*-OE Plants Compared to WT Plants.

**Additional file 12: Data Set 3.** Differentially expressed genes in *OsLG3*-RNAi Plants Compared to WT Plants.

